# Proresolving mediators LXB_4_ and RvE1 regulate inflammation in stromal cells from patients with shoulder tendon tears

**DOI:** 10.1101/606152

**Authors:** Stephanie G Dakin, Romain A Colas, Kim Wheway, Bridget Watkins, Louise Appleton, Jonathan Rees, Stephen Gwilym, Christopher Little, Jesmond Dalli, Andrew J Carr

## Abstract

Tendon stromal cells isolated from patients with chronic shoulder rotator-cuff tendon tears show dysregulated resolution responses. Current therapies do not address the biological processes concerned with persistent tendon inflammation, therefore new therapeutic approaches targeting tendon stromal cells are required. We determined if two specialised pro-resolving mediators (SPM) LXB_4_ and RvE1, modulated the bioactive lipid mediator (LM) profiles of IL-1β stimulated tendon cells derived from patients with shoulder tendon tears and healthy volunteers. We also determined if LXB_4_/RvE1 treatments moderated the pro-inflammatory phenotype of tendon tear stromal cells. Incubation of IL-1β treated patient derived tendon cells in LXB_4_/RvE1 upregulated concentrations of SPM. RvE1 treatment specifically increased 15-epi-LXB_4_ and regulated PGF_2α_. LXB_4_ or RvE1 also induced expression of the SPM biosynthetic enzymes 12-liopxygeanse (ALOX12), and ALOX15. RvE1 treatment upregulated proresolving receptor ERV1 compared to vehicle treated cells. Incubation in LXB_4_ or RvE1 moderated the proinflammatory phenotype of patient derived tendon tear cells, regulating markers of tendon inflammation, including Podoplanin, CD90, STAT-1 and IL-6. These treatments also suppressed JNK1/2/3, Lyn, STAT-3 and STAT-6 and induced p70s6kinase phospho-kinase signalling. LXB_4_ and RvE1 counter-regulate inflammatory processes in tendon stromal cells, supporting the role of these molecules as potential therapeutics to resolve tendon inflammation.

## INTRODUCTION

Diseases of the joint are a considerable global economic burden, accounting for 5 of the top 15 causes of years lived with disability in well-resourced health systems^1^. Shoulder rotator cuff tendon tears are a progressive inflammatory and fibrotic condition affecting 15% of 60-year olds and 50% of 80-year olds^2, 3^ Affected patients experience pain and restricted joint motion, severely limiting activities and disrupting life quality^4^. Current treatments include physical therapy, non-steroidal anti-inflammatory drugs, platelet rich plasma, glucocorticoid injections and surgery to repair torn tendons. These therapies are frequently ineffective, glucocorticoids are potentially harmful and tendon tear surgery is associated with high post-operative failure rates^5–7^. Of importance, COX-2 selective NSAIDs dampen protective responses regulating resolution of inflammation^8, 9^, paradoxically impeding the ability of inflamed tendons to heal. To address this unmet clinical requirement, effective new therapies are required that target the biological mechanisms and cells driving tendon disease.

Growing evidence supports the pivotal role of resident stromal cells including fibroblasts in inflammatory diseases of the joint. Fibroblasts are implicated in the switch from acute to chronic inflammation^10^ Exposure to an inflammatory milieu induces fibroblasts to undergo phenotypic change whereby these cells exhibit characteristics of an activated state and show capacity for inflammation memory^11, 12^. Cross-talk between fibroblasts with tissue resident macrophages, infiltrating immune cells and endothelial cells via cytokine and chemokine gradients in inflamed tissues of the joint further promotes the development of persistent inflammation^13, 14^ We recently identified tendon stromal cells isolated from patients with shoulder tendon tears exhibited a pro-inflammatory phenotype, and showed dysregulated resolution responses compared to respective cells isolated from the tendons of healthy volunteers^15^. Specialised proresolving mediators (SPM) including 15-epi-LXA_4_ and MaR1 were found to counter-regulate the dysregulated resolution responses of diseased tendon stromal cells^15^ This study also identified SPM including Lipoxin B_4_ (LXB_4_) and E series resolvins were differentially regulated in cultures of tendon stromal cells isolated from patients with shoulder tendon tears compared to cells from the tendons of healthy volunteers^15^. The main objective of the current study was to identify new therapeutic approaches to target pathogenic stromal cells and promote resolution of inflammation in cells isolated from patients with shoulder tendon tears. Through experiments with representative SPM including LXB_4_ and RvE1, we provide evidence that these SPM regulate the proinflammatory phenotype and promote resolution responses in patient-derived tendon stromal cells.

## MATERIALS AND METHODS

### Study approval

Tendon tissues were collected from patients under Research Ethics from the Oxford Musculoskeletal Biobank (09/H0606/11). Full informed consent according to the Declaration of Helsinki was obtained from all patients.

### Collection of patient tendon tissues

Patients with rotator cuff shoulder tendon tears were recruited from orthopaedic referral clinics. Patients had failed non-operative treatment, including a course of physical therapy, and had experienced pain for a minimum of 3 months. The presence of a supraspinatus tendon tear was identified by ultrasound scan. Patients completed the Oxford Shoulder Score (OSS), a validated and widely used clinical outcome measure scoring from 0 (severe pathology) to 48 (normal function). Supraspinatus tendon tears were collected at the time of surgical debridement of the edges of the torn tendons from 15 male and female patients aged between 46 and 75 (mean 57 ± 16.3 years). All patients were symptomatic and had small to medium tears (≤1 cm to ≤3 cm in anterior-posterior length). Exclusion criteria for all patients in this study included previous shoulder surgery, other shoulder pathology and inflammatory arthritis. Diabetic patients and those receiving systemic anticoagulant therapy were also excluded from the study. Samples of healthy volunteer hamstring tendons were collected from 10 male and female patients undergoing surgical reconstruction of their anterior cruciate ligament. All healthy volunteer patients were aged between 20 and 45 (mean 27.2 ± 10 years).

### Isolation of tendon-derived stromal cells from healthy and diseased tendons

Tendon derived stromal cells were isolated from the tendons of patients and healthy volunteers using previously published protocols^12, 16^ For experiments, cells were incubated in DMEM F12 media (Gibco) containing 1% heat inactivated human serum (Sigma) and 1% Pen-Strep. Passage 1–3 cells were used for all experiments. We previously characterised tendon stromal cells as CD45^neg^ and CD34^neg cells^ exhibiting fibroblast morphology^12^.

### Cytokine treatment of tendon-derived stromal cells

IL-1β is known to induce expression of NF-κB target genes highly expressed in shoulder tendon disease^16^. We therefore investigated the bioactive LM profiles in tendon-derived stromal cells isolated from the tendons of healthy volunteers and patients in the presence of IL-1β (10 ngmL^-1^, Sigma) in medium (DMEM F12 phenol red free medium, Gibco), containing 1% heat-inactivated human serum (Sigma) and 1% penicillin-streptomycin. Non-treated (vehicle only) cells served as experimental controls. After cytokine/vehicle treatment, cells were incubated at 37°C and 5% CO_2_ for 24 hours until experimental harvest of the media and lysate for bioactive LM profiling.

### Modulating bioactive lipid mediator profiles of IL-1β stimulated tendon-derived stromal cells with LXB_4_ and RvE1

Tendon stromal cells were isolated from healthy volunteers or tendon tear patients (n-5 each) and seeded at a density of 60,000 cells per well. Once cells were 80% confluent, they were pre-incubated with 10 nM LXB_4_ (Cayman Chemical) or 10 nM RvE1 (Cayman Chemical) for 24 h in DMEM F12 phenol red free medium (Gibco) containing 1% heat inactivated human serum (Sigma) and 1% penicillin-streptomycin. Cells were stimulated with IL-1β (10ngml^-1^) in the presence of media containing either LXB_4_, RvE1 or vehicle control as previously described^15^ Parallel experiments were performed and cell lysates harvested to investigate if incubating cells in these SPM moderated the expression of markers of the pro-inflammatory phenotype of diseased tendon stromal cells and potentiated expression of SPM synthetic enzymes and receptors mediating resolution of inflammation. The concentration and integrity of mediators used for these incubations were validated using UV-spectrophotometry and LC-MS-MS in accordance with published criteria^17^. Bioactive LM profiling of media and lysate samples from IL-1β stimulated tendon cells was performed using previously described methodology^15^ Calibration curves were obtained for each using authentic compound mixtures and deuterium labelled LM at 0.78, 1.56, 3.12, 6.25, 12.5, 25, 50, 100, and 200 pg. Linear calibration curves were obtained for each LM, which gave r^2^ values of 0.98–0.99.

### Immunocytochemistry for LXB_4_ and RvE1 treated tendon stromal cells

Tendon stromal cells isolated from patients and healthy volunteers were grown in chamber slides and stimulated with IL-1β in the presence of LXB_4_, RvE1 or vehicle for 24 hrs as described above. Cells were fixed in ice cold methanol for 5 mins and washed with PBS. Immunofluorescence staining protocols and image acquisition are adapted from a previously published protocol^15^. Tendon stromal cells isolated from healthy volunteers and patients with tendon tears (n=3 each) were incubated with the following primary antibodies: anti-ALX (Abcam, ab26316), anti-ALOX15 (Abcam ab119774), anti-ALOX12 (Abcam, ab211506), anti-ERV1 (Abcam ab167097), anti-BLT1 (Abcam ab18886), anti-STAT-1 (Abcam ab29045), anti-Podoplanin (Abcam ab10288) and anti-IL-6 (Abcam ab9324) in PBS containing 5% goat serum in Saponin for 3 hrs at room temperature. For negative controls the primary antibody was substituted for universal isotype control antibodies: cocktail of mouse IgG1, IgG_2a_, IgG_2_b, IgG_3_ and IgM (Dako) and rabbit immunoglobulin fraction of serum from non-immunized rabbits, solid phase absorbed. Isotype control staining is shown in Figure S1. Immunofluorescence images were acquired on a Zeiss LSM 710 confocal microscope using a previously published protocol^15^.

### Expression of proinflammatory and proresolving genes in tendon stromal cells incubated in LXB_4_ or RvE1

Tendon-derived stromal cells from healthy volunteers or patients with shoulder tendon tears (n=6 each) were seeded at a density of 20,000 cells per well in a 24 well plate. Cells were allowed to reach confluence prior to preincubation with LXB_4_ or RvE1 and subsequent stimulation with IL-1β (10ngmL^-1^). Non-treated cells (vehicle only, containing 0.1% endotoxin free BSA, Sigma) served as controls for each experiment. After treatment, cells were then incubated at 37 °C and 5% CO_2_ until harvest of the cell lysate for mRNA after 24 h. RNA isolation, cDNA synthesis and quantitative PCR were performed using previously published protocols^16^. Pre-validated Qiagen primer assays (*ALOX15, ERV1, IL6, PDPN, CD90, β-actin* and *GAPDH)* were used for qPCR. Results were calculated using the ddCt method using reference genes for human *β-actin* and *GAPDH*. Results were consistent using these reference genes and data are shown normalized to *β-actin*.

### Quantification of Interleukin-6 in tissue culture media

IL-6 is an important cytokine implicated in inflammation and is abundantly released by tendon stromal cells isolated from patients with shoulder tendon tears after stimulation with IL-1β^15^ IL-6 in tissue culture supernatants was measured using enzyme-linked immunosorbent assay (ELISA) reagents (BD Biosciences) using incubations isolated from 5 donors. Minimum detectable IL-6 concentration for this assay was 2.2 pgml^-1^. Optical density was read on a spectrophotometric ELISA plate reader (FLUOstar Omega, BMG Labtech) and analysed using MARS data analysis software.

### Phospho-signalling in LXB_4_ and RvE1 treated tendon stromal cells

A human phospho-kinase array kit (R&D Systems ARY003B) was used to investigate the effects of incubating IL-1β stimulated patient derived tendon cells in LXB_4_ or RvE1 on protein kinase signalling pathways (n=3 donors). Experimental protocols were performed according to manufacturer’s instructions on protein lysates harvested after 24h incubation in either LXB_4_ or RvE1. Images were captured using a chemiluminescence documentation system (UVITEC), and densitometry analysis of proteins of interest was performed using ImageJ software (NIH).

### Statistical analysis

Statistical analyses were performed using GraphPad Prism 7 (GraphPad Software). Normality was tested using the Shapiro-Wilk normality test. Analysis of bioactive LM profiles from tendon cells derived from patients and healthy volunteers was performed using multivariate statistical analysis, orthogonal-partial least squares-discriminant analysis (o-PLS-DA) using SIMCA 14.1 software (Umetrics, Umea, Sweden) following unit variance scaling of LM amounts. PLS-DA is based on a linear multivariate model that identifies variables that contribute to class separation of observations (cell incubations) on the basis of their variables (LM levels). During LM classification, observations were projected onto their respective class model. The score plot illustrates the systematic clusters among the observations (closer plots presenting higher similarity in the data matrix). Loading plot interpretation identified the variables with the best discriminatory power (Variable Importance in Projection greater then 1) that were associated with tight clusters for LM profiles obtained from incubations with cells from healthy volunteers or patients with tendinopathy. For levels of proresolving mediators and inflammation initiating eicosanoids, data are shown as summed with SEM, where n is the biological replicate. Unpaired t-tests were used to test for differences in LM levels between tendon cells derived from healthy volunteers and patients with shoulder tendon tears. Pairwise Mann Whitney U tests were used to determine differences in expression of proinflammatory and proresolving genes and IL-6 protein in IL-1β treated tendon stromal cells in the presence or absence of LXB_4_, RvE1 or respective vehicle. *P* < 0.05 was considered statistically significant.

## RESULTS

### LXB_4_ and RvE1 treatments induce SPM release from tendon-derived stromal cells

We previously identified that tendon stromal cells isolated from patients with shoulder tendon tears show dysregulated resolution responses compared to cells isolated from healthy volunteer tendons^15^. This study identified SPM including LXB_4_ and E series resolvins were differentially regulated in these incubations. To gain further insights into whether these SPM counter-regulate tendon inflammation, we therefore investigated whether LXB_4_ and RvE1 modulated the bioactive LM profiles of IL-1β stimulated tendon stromal cells isolated from the tendons of patients with shoulder tendon tears and healthy volunteers. Multivariate analysis identified differences in bioactive LM profiles between IL-1β stimulated tendon cells isolated from either healthy volunteer donors or patients with tendon tears in the presence of 10nM LXB_4_ compared to vehicle only incubations, demonstrated by the distinct clustering of the LM profiles (Figure 1A-D). The molecules profiled, together with the concentrations of the individual lipid mediators identified are listed in Table S1. In these incubations, LXB_4_ upregulated concentrations of SPM in IL-1β stimulated tendon cells from healthy volunteers (p=0.008) and tendon tear patients (p=0.008) (Figure 1E). Multivariate analysis also identified distinct clustering of the LM profiles between IL-1β stimulated tendon cells isolated from healthy volunteer donors and tendon tear patients incubated in the presence of 10nM RvE1 compared to vehicle only (Figure 2A-D). The molecules profiled, together with the concentrations of the individual lipid mediators identified are listed in Table S1. In these incubations, RvE1 upregulated concentrations of SPM in healthy volunteer tendon cells (p=0.008) and tendon tear patients (p=0.008) (Figure 2E). In tendon tear incubations, RvE1 upregulated the concentrations of specific SPM including 15-epi-LXB_4_ (p=0.04) and decreased levels of the pro-inflammatory eicosanoid PGF_2α_ (p=0.02) (Figure 2E).

**FIGURE 1.**
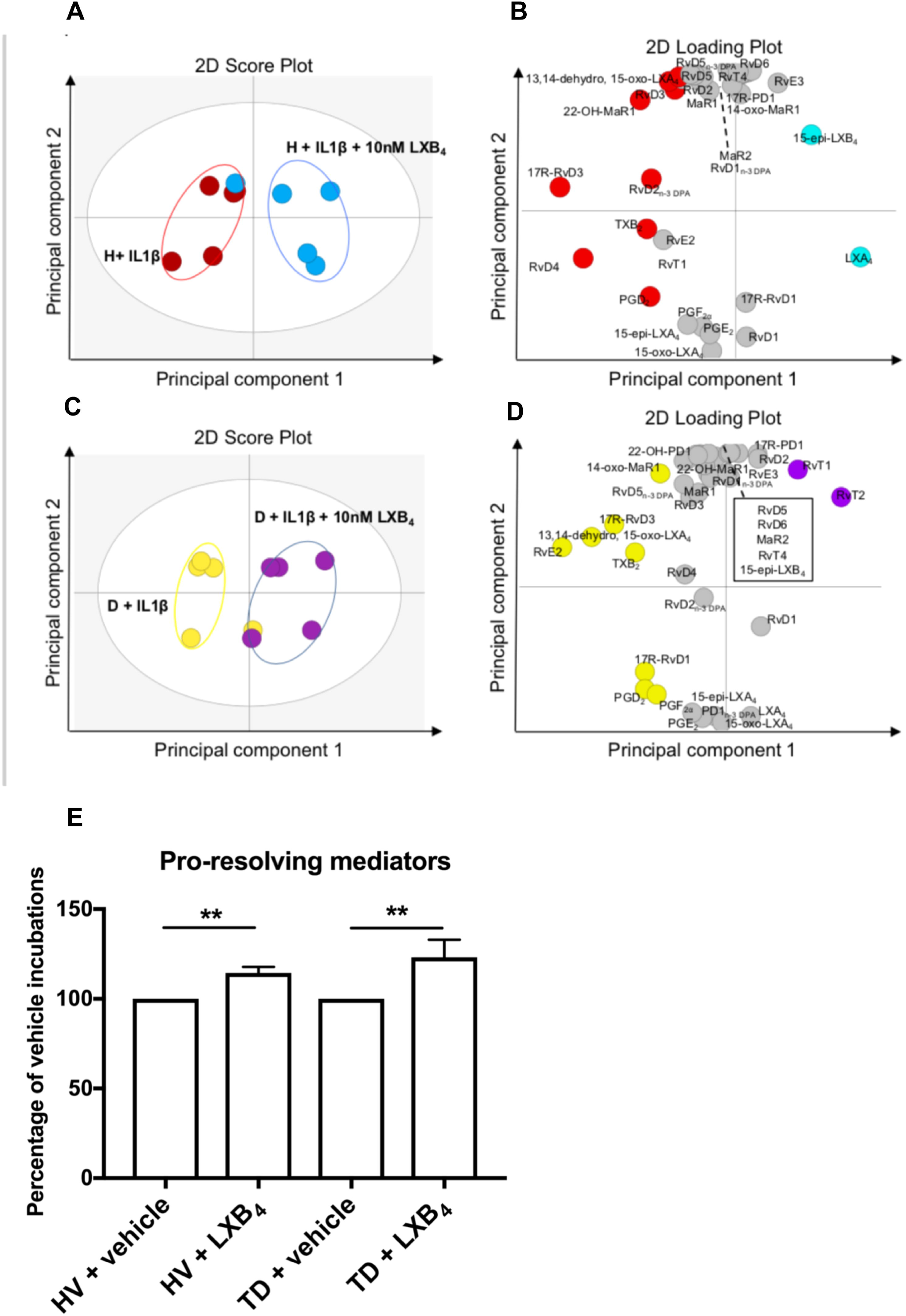
LXB_4_ upregulates SPM concentrations in IL-1β stimulated tendon stromal cells. Tendon stromal cells were derived from healthy volunteers (Healthy, H, n=5 donors) and patients with shoulder tendon tears (Diseased, D n=5 donors). Cells were incubated with LXB_4_ (10nM) or vehicle for 24 h at 37 °C then with IL-1β (10ngml^-1^) for 24 h. LM were identified and quantified using LM profiling. **(A)** 2-dimensional score plot and **(B)** corresponding 2-dimensional loading plot of LM-SPM from human tendon derived-stromal cell incubations isolated from healthy volunteers incubated with IL-1β and 10nM LXB_4_ or vehicle only. **(C)** 2-dimensional score plot and **(D)** corresponding 2-dimensional loading plot of LM-SPM from human tendon derived-stromal cell incubations isolated from patients with shoulder tendon tears incubated with IL-1β and 10nM LXB_4_ or vehicle only. **(E)** Cumulative concentrations of proresolving mediators (DHA-derived RvD, PD, MaR, n-3 DPA-derived RvD_n-3 DPA_, PD_n-3 DPA_, MaR_n-3 DPA_, EPA-derived RvE and AA-derived LX) in IL-1β stimulated tendon stromal cell incubations in the presence of LXB_4_ (10nM) or vehicle for 24 h. Results are shown as means and SEM and representative of n=5 donors per group.

**FIGURE 2.**
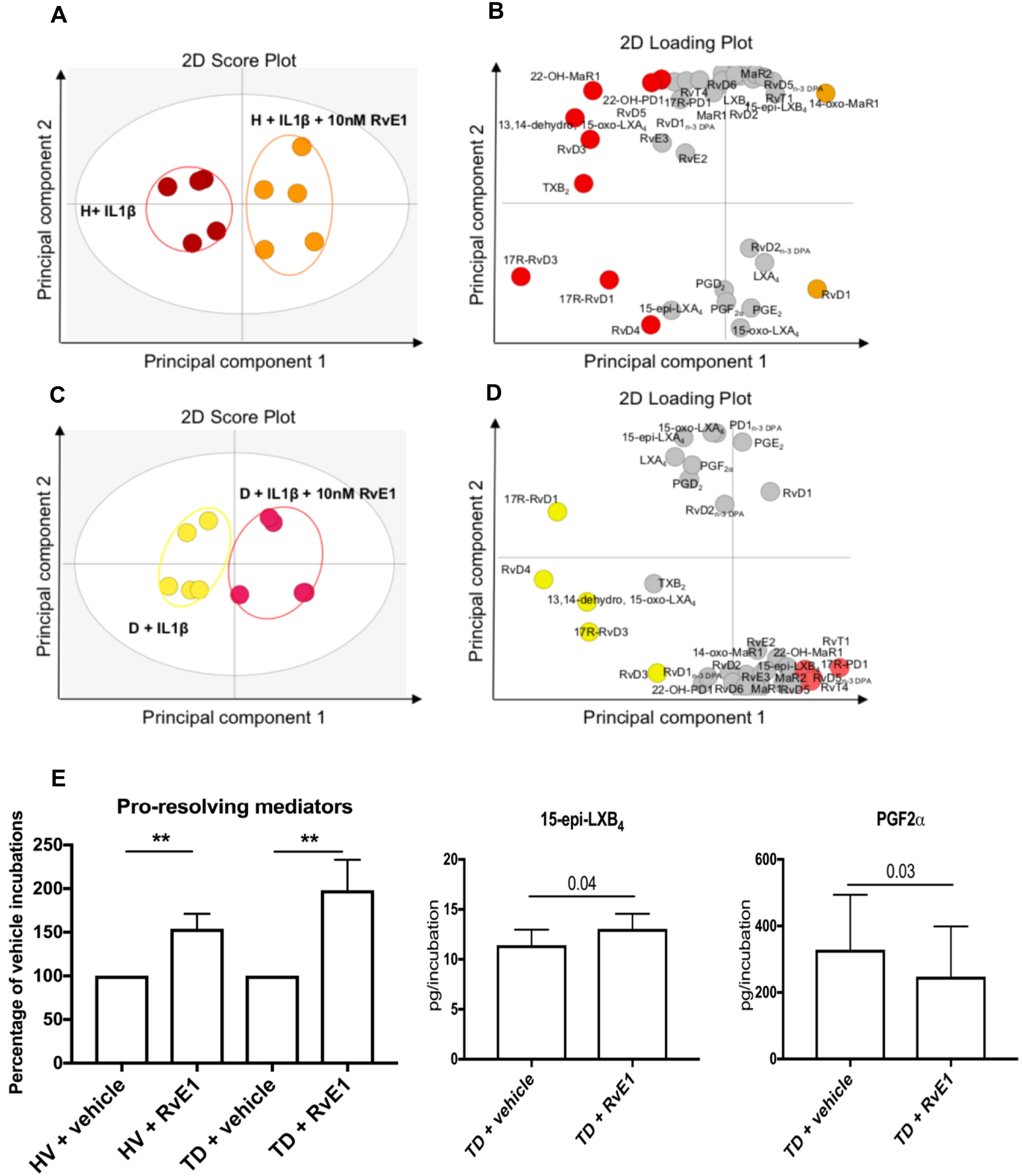
RvE1 increases SPM levels in IL-1β stimulated tendon stromal cells. Tendon stromal cells were derived from healthy volunteers (Healthy, H, n=5 donors) and patients with shoulder tendon tears (Diseased, D n=5 donors). Cells were incubated with RvE1 (10nM) or vehicle for 24 h at 37 °C then with IL-1β (10ngml^-1^) for 24 h. LM were identified and quantified using LM profiling. **(A)** 2-dimensional score plot and **(B)** corresponding 2-dimensional loading plot of LM-SPM from human tendon derived-stromal cell incubations isolated from healthy volunteers incubated with IL-1β and 10nM RvE1 or vehicle only. **(C)** 2-dimensional score plot and **(D)** corresponding 2-dimensional loading plot of LM-SPM from human tendon derived-stromal cell incubations isolated from patients with shoulder tendon tears incubated with IL-1β and 10nM RvE1 or vehicle only. **(E)** Cumulative levels of proresolving mediators and differentially regulated lipid mediators in IL-1β stimulated tendon stromal cell incubations in the presence of RvE1(10nM) or vehicle for 24 h. Results are shown as means and SEM and representative of n=5 donors per group.

### LXB_4_ and RvE1 upregulate the expression of SPM biosynthetic enzymes and proresolving receptors in tendon-derived stromal cells

We next investigated the mechanisms by which LXB_4_ and RvE1 potentiated the further release of SPM. Incubation of IL-1β stimulated tendon cells isolated from tendon tear patients in LXB_4_ or RvE1 induced *ALOX15* mRNA expression relative to respective vehicle controls (p=0.03 respectively, Figure 3A). The same treatment of healthy volunteer tendon cells also upregulated *ALOX15* mRNA expression relative to respective vehicle controls (p=0.03 and p=0.01 respectively, Figure 3A). Immunostaining demonstrated these treatments also increased ALOX12 and ALOX15 proteins expression implicated in the biosynthesis of SPM (Figure 3B and 3C). Induction of these biosynthetic enzymes was profound in tendon cells isolated from tendon tear patients compared to healthy volunteer donors (Figures 3B&C). We also investigated if incubation of tendon tear cells in RvE1 moderated expression of receptors to which RvE1 is known to bind. In the presence of vehicle only, IL-1β stimulated tendon tear cells showed increased *CHEMR23/ERV1*mRNA compared to respective healthy tendon cells (Figure 3D). Indeed, RvE1 treatment upregulated the expression of ChemR23/ERV1 and BLT1 receptors in IL-1β stimulated tendon tear cells compared to respective vehicle controls (Figure 3E).

**FIGURE 3.**
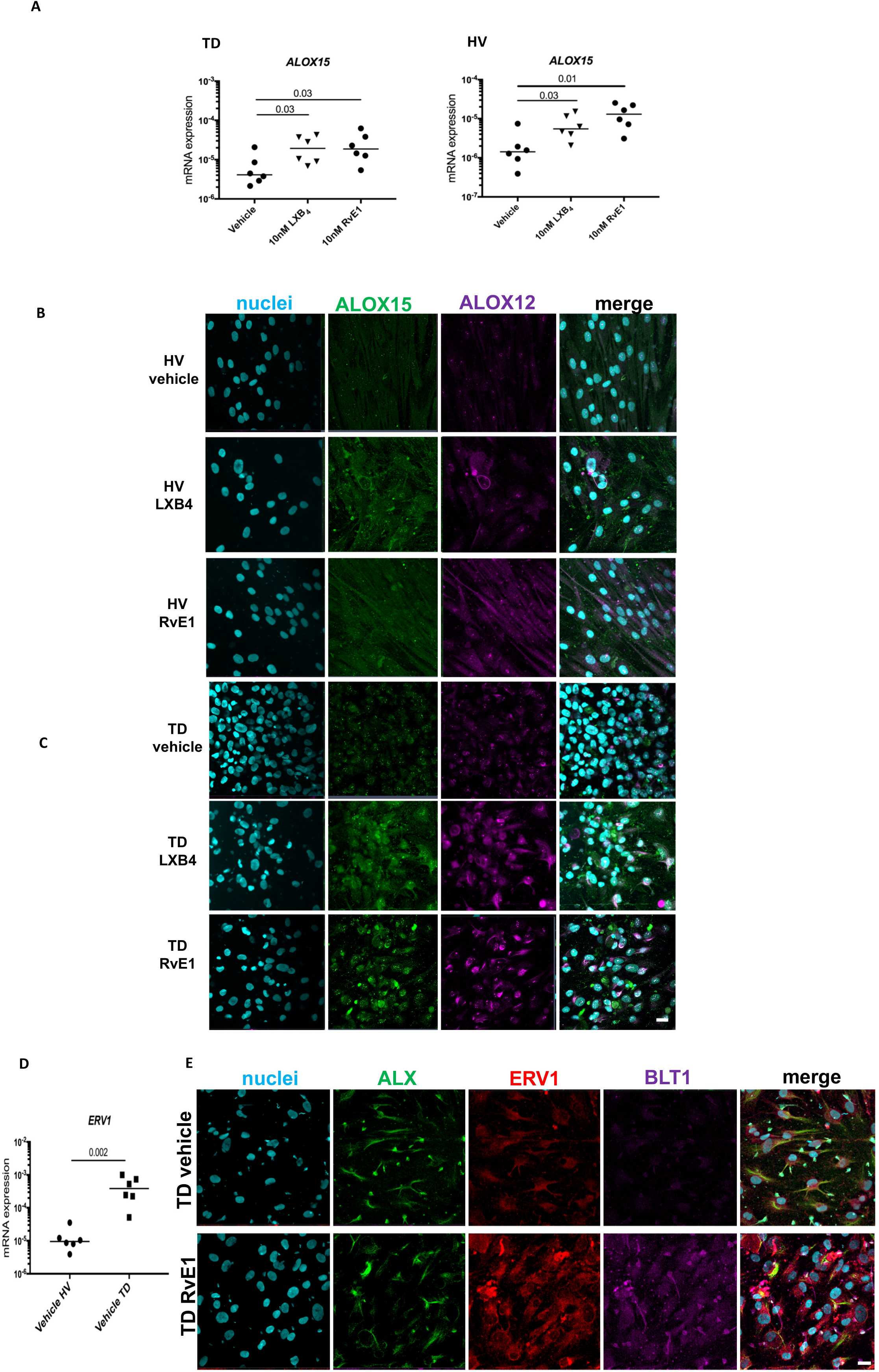
LXB_4_ and RvE1 induce SPM biosynthetic enzymes and regulate the pro-resolving receptor ChemR23/ERV1 in tendon stromal cells. Tendon stromal cells were derived from patients with shoulder tendon tears (TD, n=6) or healthy volunteers (HV, n=6). Cells were incubated with LXB_4_ (10nM), RvE1 (10nM) or vehicle for 24 h at 37 °C then with IL-1β (10ngml^-1^) for 24 h. **(A)** Incubation in LXB_4_ significantly induced *ALOX15* mRNA in both TD and HV cells (p=0.03 respectively) compared to respective vehicle controls. Incubation in RvE1 significantly induced *ALOX15* mRNA in both TD (p=0.03) and HV cells (p=0.01) compared to respective vehicle controls. Gene expression is normalized to β-actin; bars show median values. Representative images of immunocytochemistry for SPM synthetic enzymes ALOX15 (green) ALOX12 (violet) in IL-1β stimulated HV **(B)** and TD tendon stromal cells **(C)** incubated in 10nM LXB_4_, 10nM RvE1 or vehicle control for 24 hrs. Cyan represents POPO-1 nuclear counterstain. All images are representative of n=3 donors. Scale bar, 20 μm. **(D)** *ERV1* mRNA expression in IL-1β stimulated HV and TD cells. Gene expression is normalized to β-actin; bars show median values. **(E)** Representative images of immunocytochemistry for proresolving receptors ALX (green), ERV1 (red) and BLT1 (violet) in IL-1β stimulated diseased tendon stromal cells incubated in 10nM LXB_4_, 10nM RvE1 or vehicle control for 24 hrs. Cyan represents POPO-1 nuclear counterstain. All images are representative of n=3 donors. Scale bar, 20 μm.

### LXB_4_ and RvE1 moderate the pro-inflammatory phenotype of tendon stromal cells, dampening pro-inflammatory signalling pathways

We next assessed whether LXB_4_ and RvE1 also regulated known markers of tendon inflammation in tendon cells isolated from tendon tear patients and healthy volunteers. Incubation of IL-1β stimulated diseased cells in LXB_4_ or RvE1 for 24 hrs reduced fibroblast activation marker Podoplanin (PDPN), STAT-1 and IL-6 compared to respective vehicle controls (Figure 4A). The same treatment of IL-1β stimulated healthy tendon cells also reduced PDPN, STAT-1 and IL-6 compared to respective vehicle controls (Figure 4B). Measurement of IL-6 levels in supernatant from IL-1β stimulated diseased cells demonstrated incubation in LXB_4_ or RvE1 reduced IL-6 levels compared to vehicle only (p=0.02 and 0.006 respectively, Figure 4C). In HV incubations, LXB_4_ or RvE1 treatment also reduced IL-6 levels (p=0.03 and p=0.01 respectively, Figure 4D). We next determined if LXB_4_ or RvE1 treatment moderated expression of pro-inflammatory genes and signalling pathways in IL-1β stimulated tendon tear cells. Incubation of these cells in LXB_4_ reduced *IL6, PDPN* and *CD90* mRNA expression compared to vehicle controls (p=0.004, p=0.002 and p=0.015 respectively, Figure 4E). RvE1 treatment of IL-1β stimulated tendon tear cells also reduced *IL6, PDPN* and *CD90* mRNA compared to vehicle controls (p=0.02, p=0.02 and p=0.015 respectively, Figure 4E). In these incubations, LXB_4_ or RvE1 treatments regulated phospho-kinase signalling pathways identified in inflamed tendons, including JNK1/2/3 (phosphorylation sites T183/Y185, T221/Y223), Lyn (Y397), STAT-3 (Y705) and STAT-6 (Y641) compared to respective vehicle control treated cells (Figure 5). Incubation in RvE1 also induced p70S6 (T389) kinase compared to respective vehicle controls (Figure 5).

**FIGURE 4.**
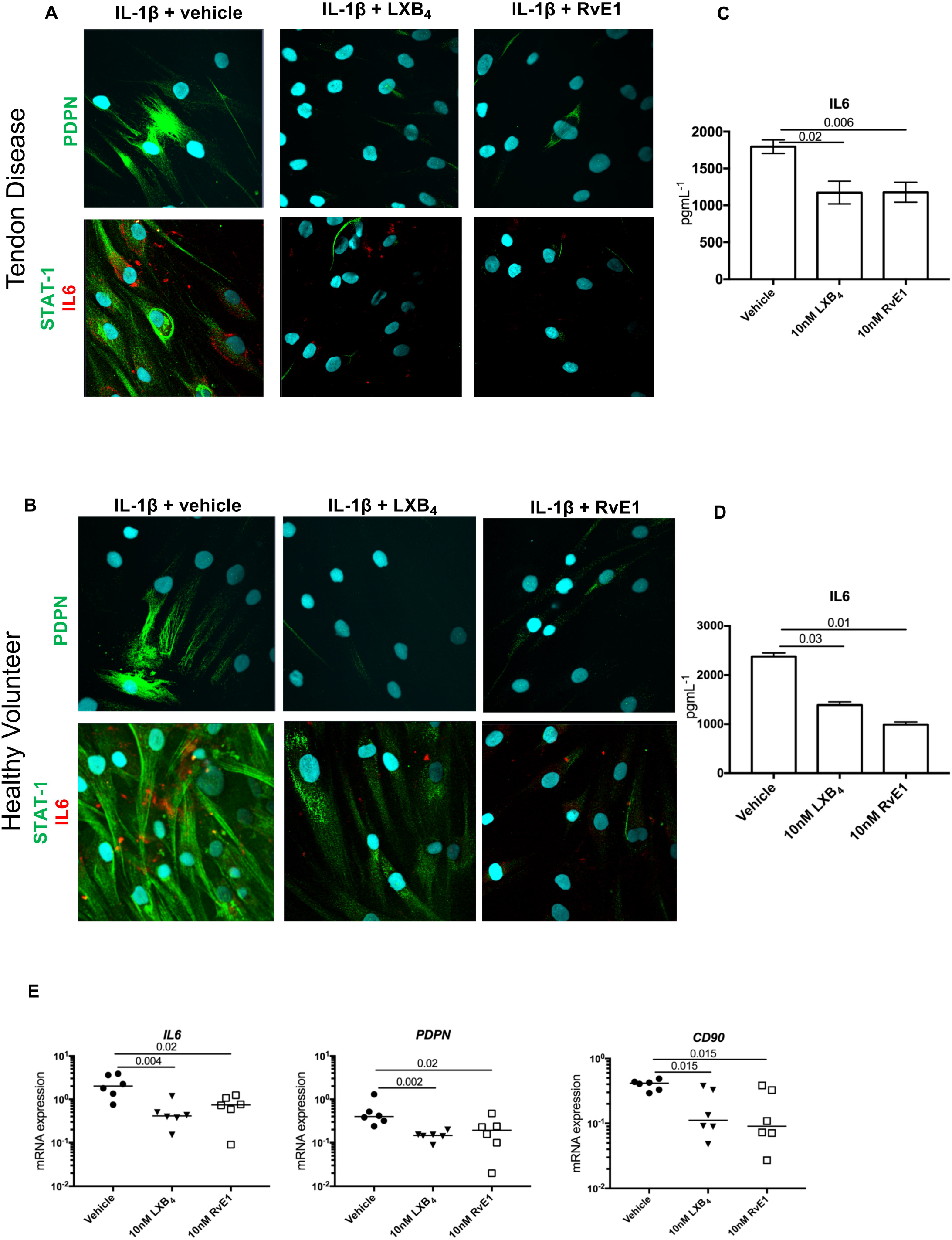
LXB_4_ and RvE1 moderate the pro-inflammatory phenotype of tendon stromal cells. Tendon stromal cells were derived from patients with shoulder tendon tears (TD) or healthy volunteers (HV). Representative images of immunocytochemistry for established markers of tendon inflammation including Podoplanin (PDPN) green, STAT-1 (green) and Interleukin-6 (IL-6, red) in IL-1β stimulated **(A)** TD and **(B)** HV tendon stromal cells incubated in 10nM LXB_4_, 10nM RvE1 or vehicle control for 24 hrs. Cyan represents POPO-1 nuclear counterstain. All images are representative of n=3 donors. Scale bar, 20μm. ELISA assay of IL-6 protein secretion from IL-1β stimulated TD **(C)** and HV **(D)** tendon cells incubated in the presence and absence of 10 nM LXB_4_ or 10nM RvE1. Data are shown as means and SEM, n=5 separate donors. **(E)** mRNA expression of markers of tendon inflammation including *IL-6,* and fibroblast activation markers PDPN and *CD90*, in IL-1β stimulated TD cells (n=6 donors) incubated in either 10nM LXB_4_, 10nM RvE1 or vehicle control for 24 hrs. Gene expression is normalized to β-actin, bars show median values.

**Figure 5.**
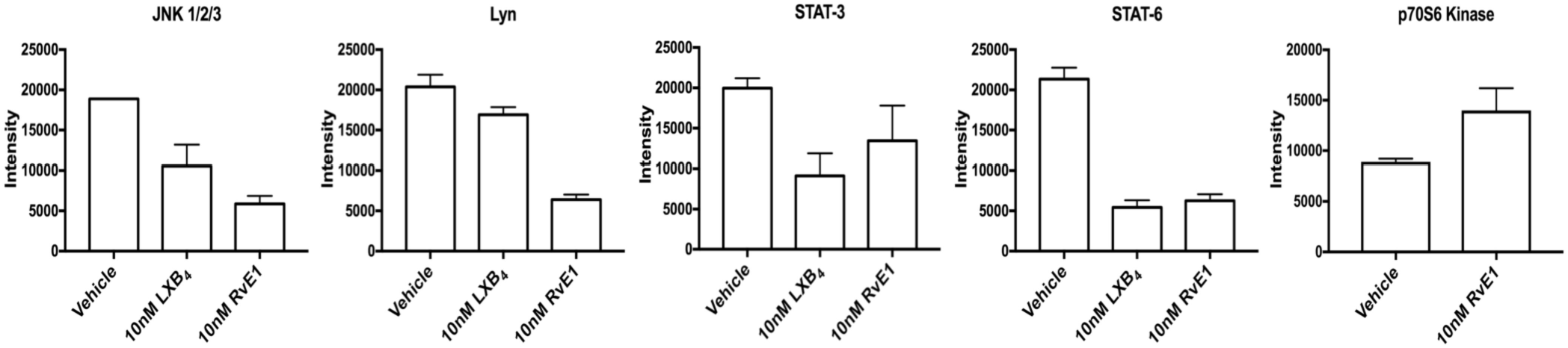
LXB_4_ and RvE1 moderate protein kinase expression in diseased tendon stromal cells. Densitometric analysis was acquired using Image J software to identify the effects of incubating IL-1β treated TD cells in LXB_4_ or RvE1 on protein phospho-kinase signalling pathways JNK1/2/3, Lyn, STAT3, STAT6 and p70s6kinase. Results are shown as means and SEM and representative of n=3 donors per group relative to respective vehicle control treated cells.

## DISCUSSION

Resident stromal cells including fibroblasts are increasingly recognised as important cell types driving chronic inflammatory joint disease^10, 18, 19^. After exposure to an inflammatory milieu, tendon and synovial fibroblasts adopt a pro-inflammatory phenotype exhibiting activation and inflammation memory^11, 12, 20^ Distinct fibroblast subtypes that mediate joint inflammation and tissue damage have been characterised in rheumatoid synovium^21^. Recent advances in the knowledge of how resident stromal cells behave under inflammatory conditions of the joint has prompted further investigation of the resolution responses of these cells. Given that stromal fibroblasts comprise the majority cell type of musculoskeletal soft tissues, improved understanding of how these cells respond to an inflammatory milieu is required to inform the development of therapeutic strategies targeting these cells. We recently identified that tendon stromal cells isolated from patients with tendon tears showed increased levels of SPM and inflammation initiating eicosanoids compared to cells isolated from healthy volunteer tendons, reminiscent of a dysregulated resolution response characteristic of chronic inflammation^15^. We also identified that incubation of IL-1β stimulated tendon stromal cells in either 15-epi-LXA4 or MaR1 regulated pro-inflammatory eicosanoids and potentiated the further release of SPM^15^. These treatments moderated the pro-inflammatory phenotype of IL-1β stimulated diseased tendon stromal cells, dampening expression of PDPN, STAT-1 and IL-6. We previously identified that SPM including LXB_4_ and E series resolvins were differentially regulated between IL-1β stimulated tendon cells isolated from patients with shoulder tendon tears compared to cells from healthy volunteer tendons^15^. In the current study, we therefore investigated if LXB_4_ or RvE1 modulated the bioactive lipid mediator (LM) profiles of IL-1β stimulated tendon cells derived from these patient cohorts. Tendon stromal cells were stimulated with IL-1β, as this cytokine is known to induce expression of NF-κB target genes highly expressed in human tendon disease^16, 20^, simulating an inflammatory milieu. In these incubations, treatment with LXB_4_ or RvE1 upregulated SPM concentrations. RvE1 treatment specifically increased 15-epi-LXB_4_ and regulated PGF2α in incubations of IL-1β stimulated diseased tendon cells. We next investigated the mechanism of action underpinning these observations. Incubating in LXB_4_ or RvE1 induced expression of SPM biosynthetic enzymes ALOX12, and ALOX15 in healthy and diseased tendon stromal cells. Notably, expression of these SPM biosynthetic enzymes was increased in diseased compared to healthy tendon stromal cells. We previously identified that incubation of tendon stromal cells in 15-epi-LXA_4_ or MaR1 induced ALOX15 expression^15^. The findings from the current study support these observations, suggesting a common mechanism whereby proresolving mediator activate feed forward cascades leading to the upregulation of other SPM via induction of ALOX biosynthetic enzymes. We next investigated if incubating tendon cells in LXB_4_ or RvE1 influenced expression of proresolving receptors. The receptor to which LXB_4_ binds have not yet been identified, while RvE1 is known to activate ERV1 and a competitive inhibitor of BLT1^22, 23^. In the absence of SPM treatment, *ERV1* mRNA expression was increased in diseased compared to healthy IL-1β stimulated tendon stromal cells, suggesting a pro-inflammatory phenotype favours increased expression of this proresolving receptor. Incubation of LXB_4_ or RvE1 did not induce ALX expression, although RvE1 treatment further upregulated ERV1 and BLT1 expression on tendon stromal cells compared to vehicle treated cells. Collectively these findings suggest a positive feedback loop, whereby RvE1 treatment upregulates ERV1 and BLT1 receptor expression. This may occur as a direct consequence of RvE1 treatment, or via RvE1 induced upregulation SPM.

We identified that incubation in LXB_4_ or RvE1 moderated the proinflammatory phenotype of patient derived tendon tear cells, regulating known markers of tendon inflammation, including PDPN, CD90, STAT-1 and IL-6. CD90 is expressed by pathogenic synovial fibroblasts from rheumatoid arthritis patients with an inflammatory and invasive phenotype^21^. We previously identified persistent fibroblast activation may be implicated in the development of chronic tendon inflammation and increased likelihood of recurrent injury^12^. Proresolving mediators may therefore possess therapeutic utility to moderate the pro-inflammatory phenotype of tendon stromal cells via attenuating expression of pathogenic fibroblast activation markers. In support of this, other SPM including 15-epi-LXA_4_ and MaR1 also moderated PDPN expression in IL-1β stimulated diseased tendon cells^15^, suggesting this property is common to different families of SPM including lipoxins, resolvins and maresins.

In addition to moderating the pro-inflammatory phenotype of diseased tendon cells, we also identified LXB_4_ and RvE1 treatments regulated phosphokinases including pJNK1/2/3, pLyn, pSTAT-3 and pSTAT-6. The findings from our study suggest LXB_4_ and RvE1 regulate IL-6 via suppressing STAT-3 signalling in patient derived tendon stromal cells. Suppression of STAT-6 signalling may modulate IL-4 and IL-13 responsive genes known to drive fibrosis in the advanced stages of tendon disease^16, 24^. IL-6 is an important cytokine implicated in the cross talk between resident stromal cells including activated endothelial cells, tissue resident macrophages, fibroblasts and infiltrating immune cells^25–27^. Given the ability of LXB_4_ and RvE1 to regulate IL-6 release from IL-1β stimulated tendon cells, these SPM may play an important role in dampening cytokine mediated cross-talk between stromal cells which actively promotes the retention of immune cells. RvE1 treatment also induced p70s6kinase signalling in IL-1β stimulated diseased tendon cells. RvE1 is also known to activate ERV-1 signalling via rs6 phosphorylation in peripheral blood neutrophils isolated from patients with type 2 diabetes^28^. These findings suggest that circulating immune cells and resident stromal fibroblasts share common signalling pathways downstream of the ERV-1 receptor.

Therapies that promote resolution of inflammation are an important future therapeutic strategy to address pathogenic stroma in chronic inflammatory disease of the joint. The pro-resolving mediator resolvin D3 (RvD3) has been shown to regulate leukocyte infiltration pro-inflammatory eicosanoids in murine inflammatory arthritis^29^. 17R-RvD1 attenuated arthritis severity, paw oedema and leukocyte infiltration, in acute murine inflammatory arthritis^30^. The current study suggests that LXB_4_ and RvE1 regulate expression of tendon pro-inflammatory molecules including Podoplanin, CD90, STAT-1, IL-6, and dampen phosphokinases including JNK1/2/3, Lyn, STAT-3 and STAT-6. These SPM also potentiated further release of proresolving mediators in IL-1β stimulated healthy and diseased tendon cells. We therefore propose that SPM including LXB_4_ and RvE1 are potential new therapeutics to target pathogenic stromal cells and potentiate resolution of chronic tendon inflammation.

## ACKNOWLEDGEMENTS

S.G.D. is a recipient of an Oxford UCB Prize Fellowship in Biomedical Research. J.D. received funding from the European Research Council (ERC) under the European Union’s Horizon 2020 research and innovation programme (grant no: 677542) and the Barts Charity (grant no: MGU0343). J.D. is also supported by a Sir Henry Dale Fellowship jointly funded by the Wellcome Trust and the Royal Society (grant 107613/Z/15/Z). Research at NDORMS, University of Oxford is supported through the National Institute for Health Research (NIHR) Oxford Musculoskeletal Biomedical Research Centre (BRC). The views expressed are those of the authors and not necessarily those of the NHS, the NIHR of the Department of Health. We are grateful to the clinical and nursing teams at the Nuffield Orthopaedic Centre in facilitating collection of healthy and diseased tendon tissue samples used for this study.

## AUTHOR CONTRIBUTIONS

Designed research: SGD, JD, AC

Performed research: SGD, RC

Contributed reagents / analytical tools: KW, BW, LA, JR, SG, CL

Analysed data: SGD, RC, JD

Wrote the paper: SGD, RC, JD, AJC

Reviewed submitted manuscript: All authors

---

Figure S1. Isotype control staining of patient derived tendon stromal cells. Representative confocal immunofluorescence images showing merged images of stromal cells isolated from patients with shoulder tendon tears, stained with isotype control antibodies for mouse IgG_1_, IgG_2a_, IgG_2b_ and rabbit IgG fractions. Cyan represents POPO-1 nuclear counterstain. Scale bar, 20μm.

**Supplemental Table 1.**
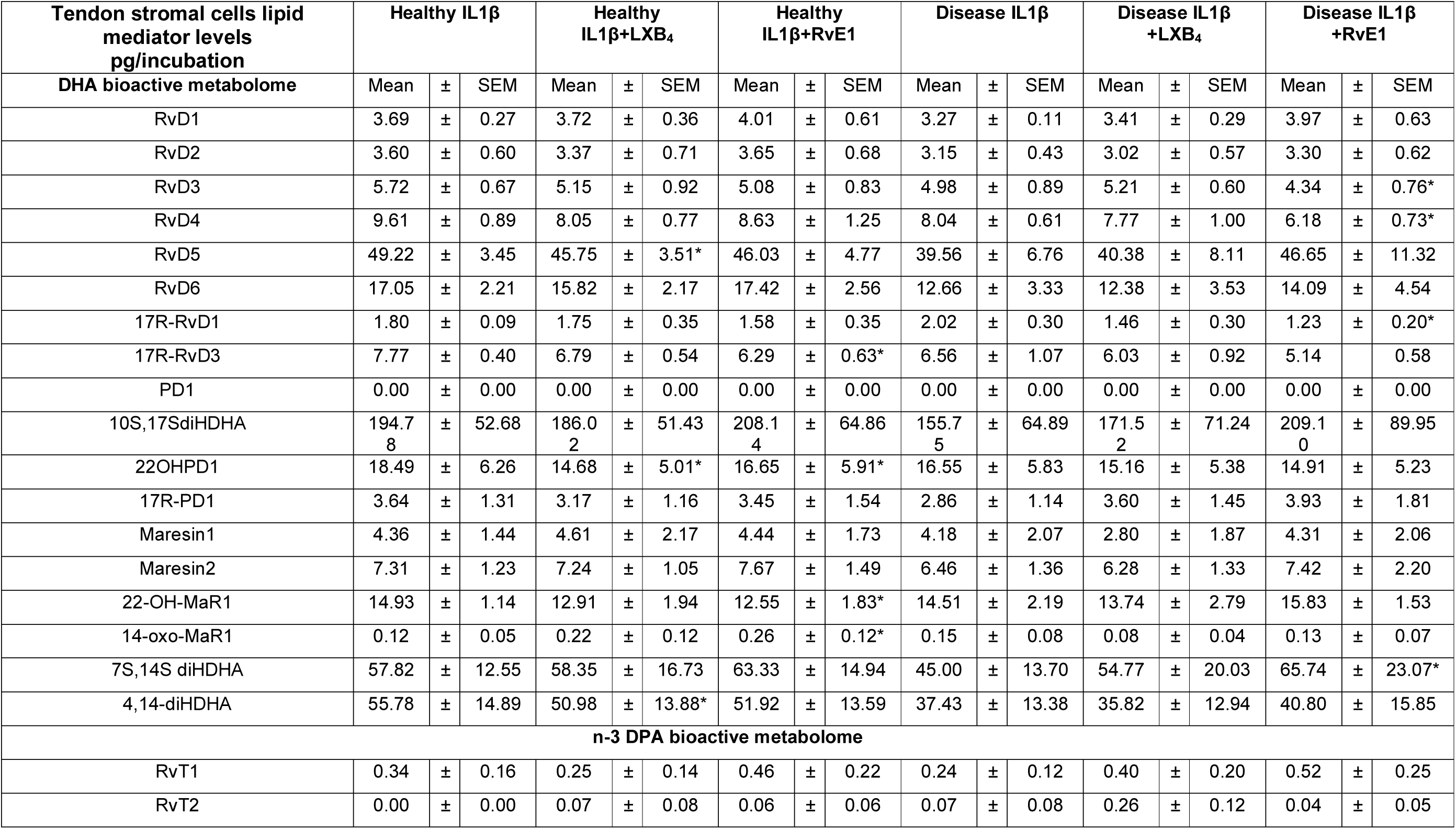
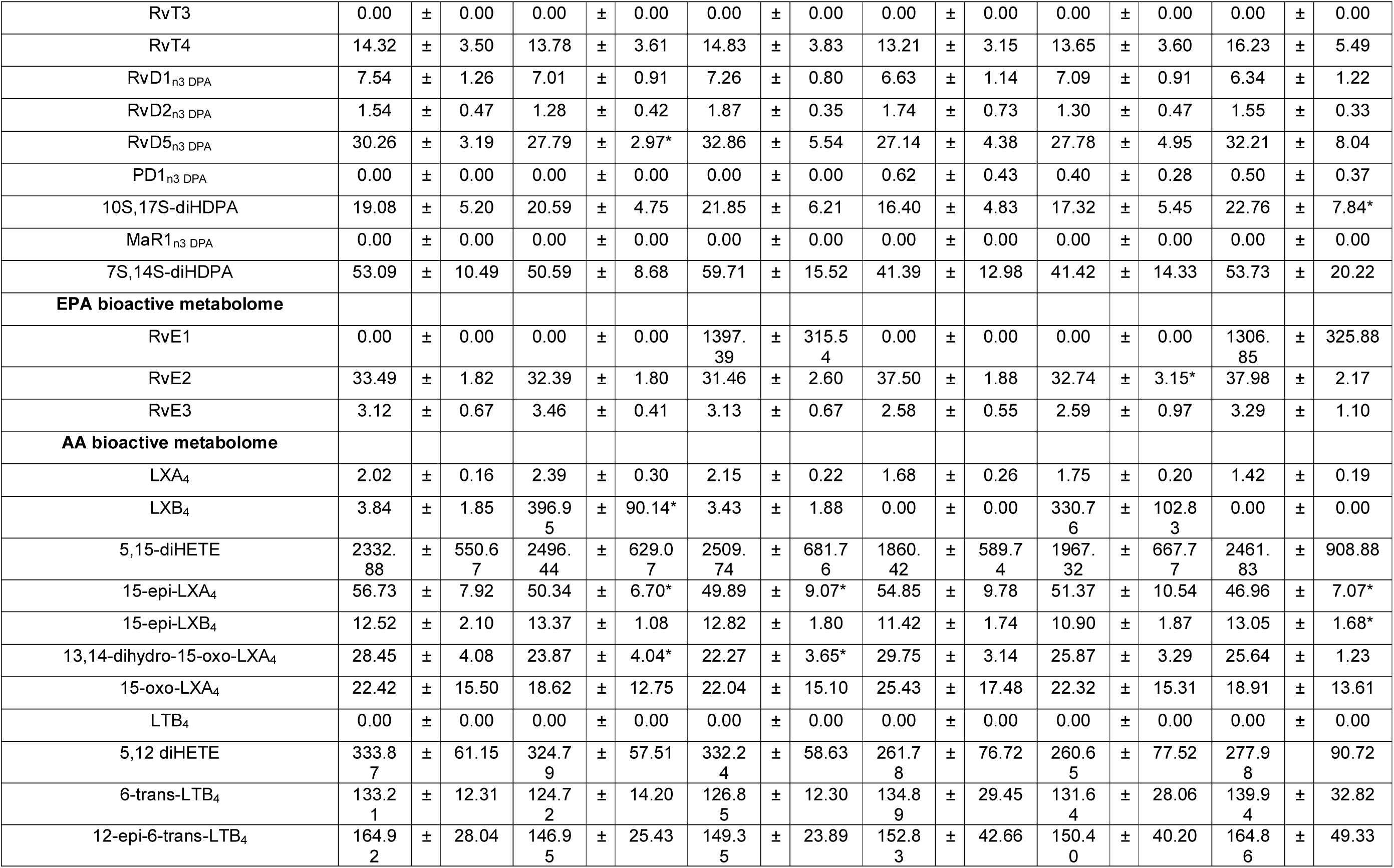
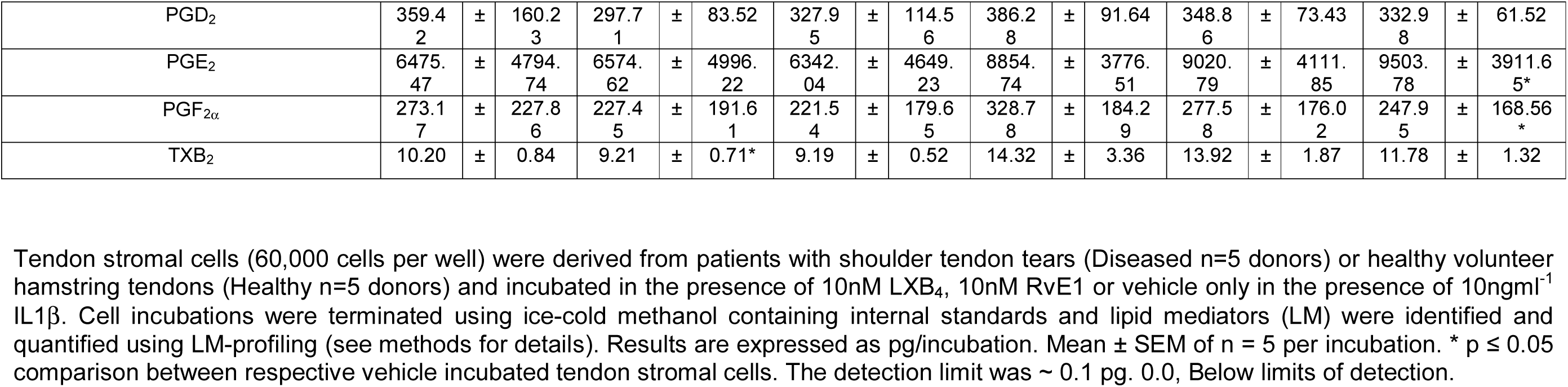
LM-SPM profiles of IL-1β stimulated patient tendon stromal cells in the presence of LXB_4_ or RvE1.

| Tendon stromal cells lipid mediator levels<br>pg/incubation | Healthy IL1β | | | Healthy IL1β+LXB<sub>4</sub> | | | Healthy IL1β+RvE1 | | | Disease IL1β | | | Disease IL1β+LXB<sub>4</sub> | | | Disease IL1β+RvE1 | | |
| --- | --- | --- | --- | --- | --- | --- | --- | --- | --- | --- | --- | --- | --- | --- | --- | --- | --- | --- |
|  | Mean | ± | SEM | Mean | ± | SEM | Mean | ± | SEM | Mean | ± | SEM | Mean | ± | SEM | Mean | ± | SEM |
| **DHA bioactive metabolome** |  |  |  |  |  |  |  |  |  |  |  |  |  |  |  |  |  |  |
| RvD1 | 3.69 | ± | 0.27 | 3.72 | ± | 0.36 | 4.01 | ± | 0.61 | 3.27 | ± | 0.11 | 3.41 | ± | 0.29 | 3.97 | ± | 0.63 |
| RvD2 | 3.60 | ± | 0.60 | 3.37 | ± | 0.71 | 3.65 | ± | 0.68 | 3.15 | ± | 0.43 | 3.02 | ± | 0.57 | 3.30 | ± | 0.62 |
| RvD3 | 5.72 | ± | 0.67 | 5.15 | ± | 0.92 | 5.08 | ± | 0.83 | 4.98 | ± | 0.89 | 5.21 | ± | 0.60 | 4.34 | ± | 0.76* |
| RvD4 | 9.61 | ± | 0.89 | 8.05 | ± | 0.77 | 8.63 | ± | 1.25 | 8.04 | ± | 0.61 | 7.77 | ± | 1.00 | 6.18 | ± | 0.73* |
| RvD5 | 49.22 | ± | 3.45 | 45.75 | ± | 3.51* | 46.03 | ± | 4.77 | 39.56 | ± | 6.76 | 40.38 | ± | 8.11 | 46.65 | ± | 11.32 |
| RvD6 | 17.05 | ± | 2.21 | 15.82 | ± | 2.17 | 17.42 | ± | 2.56 | 12.66 | ± | 3.33 | 12.38 | ± | 3.53 | 14.09 | ± | 4.54 |
| 17R-RvD1 | 1.80 | ± | 0.09 | 1.75 | ± | 0.35 | 1.58 | ± | 0.35 | 2.02 | ± | 0.30 | 1.46 | ± | 0.30 | 1.23 | ± | 0.20* |
| 17R-RvD3 | 7.77 | ± | 0.40 | 6.79 | ± | 0.54 | 6.29 | ± | 0.63* | 6.56 | ± | 1.07 | 6.03 | ± | 0.92 | 5.14 |  | 0.58 |
| PD1 | 0.00 | ± | 0.00 | 0.00 | ± | 0.00 | 0.00 | ± | 0.00 | 0.00 | ± | 0.00 | 0.00 | ± | 0.00 | 0.00 | ± | 0.00 |
| 10S,17SdiHDHA | 194.7<br>8 | ± | 52.68 | 186.0<br>2 | ± | 51.43 | 208.1<br>4 | ± | 64.86 | 155.7<br>5 | ± | 64.89 | 171.5<br>2 | ± | 71.24 | 209.1<br>0 | ± | 89.95 |
| 22OHPD1 | 18.49 | ± | 6.26 | 14.68 | ± | 5.01* | 16.65 | ± | 5.91* | 16.55 | ± | 5.83 | 15.16 | ± | 5.38 | 14.91 | ± | 5.23 |
| 17R-PD1 | 3.64 | ± | 1.31 | 3.17 | ± | 1.16 | 3.45 | ± | 1.54 | 2.86 | ± | 1.14 | 3.60 | ± | 1.45 | 3.93 | ± | 1.81 |
| Maresin1 | 4.36 | ± | 1.44 | 4.61 | ± | 2.17 | 4.44 | ± | 1.73 | 4.18 | ± | 2.07 | 2.80 | ± | 1.87 | 4.31 | ± | 2.06 |
| Maresin2 | 7.31 | ± | 1.23 | 7.24 | ± | 1.05 | 7.67 | ± | 1.49 | 6.46 | ± | 1.36 | 6.28 | ± | 1.33 | 7.42 | ± | 2.20 |
| 22-OH-MaR1 | 14.93 | ± | 1.14 | 12.91 | ± | 1.94 | 12.55 | ± | 1.83* | 14.51 | ± | 2.19 | 13.74 | ± | 2.79 | 15.83 | ± | 1.53 |
| 14-oxo-MaR1 | 0.12 | ± | 0.05 | 0.22 | ± | 0.12 | 0.26 | ± | 0.12* | 0.15 | ± | 0.08 | 0.08 | ± | 0.04 | 0.13 | ± | 0.07 |
| 7S,14S diHDHA | 57.82 | ± | 12.55 | 58.35 | ± | 16.73 | 63.33 | ± | 14.94 | 45.00 | ± | 13.70 | 54.77 | ± | 20.03 | 65.74 | ± | 23.07* |
| 4,14-diHDHA | 55.78 | ± | 14.89 | 50.98 | ± | 13.88* | 51.92 | ± | 13.59 | 37.43 | ± | 13.38 | 35.82 | ± | 12.94 | 40.80 | ± | 15.85 |
| **n-3 DPA bioactive metabolome** |  |  |  |  |  |  |  |  |  |  |  |  |  |  |  |  |  |  |
| RvT1 | 0.34 | ± | 0.16 | 0.25 | ± | 0.14 | 0.46 | ± | 0.22 | 0.24 | ± | 0.12 | 0.40 | ± | 0.20 | 0.52 | ± | 0.25 |
| RvT2 | 0.00 | ± | 0.00 | 0.07 | ± | 0.08 | 0.06 | ± | 0.06 | 0.07 | ± | 0.08 | 0.26 | ± | 0.12 | 0.04 | ± | 0.05 |
| RvT3 | 0.00 | ± | 0.00 | 0.00 | ± | 0.00 | 0.00 | ± | 0.00 | 0.00 | ± | 0.00 | 0.00 | ± | 0.00 | 0.00 | ± | 0.00 |
| RvT4 | 14.32 | ± | 3.50 | 13.78 | ± | 3.61 | 14.83 | ± | 3.83 | 13.21 | ± | 3.15 | 13.65 | ± | 3.60 | 16.23 | ± | 5.49 |
| RvD1<sub>n3</sub> DPA | 7.54 | ± | 1.26 | 7.01 | ± | 0.91 | 7.26 | ± | 0.80 | 6.63 | ± | 1.14 | 7.09 | ± | 0.91 | 6.34 | ± | 1.22 |
| RvD2<sub>n3</sub> DPA | 1.54 | ± | 0.47 | 1.28 | ± | 0.42 | 1.87 | ± | 0.35 | 1.74 | ± | 0.73 | 1.30 | ± | 0.47 | 1.55 | ± | 0.33 |
| RvD5<sub>n3</sub> DPA | 30.26 | ± | 3.19 | 27.79 | ± | 2.97* | 32.86 | ± | 5.54 | 27.14 | ± | 4.38 | 27.78 | ± | 4.95 | 32.21 | ± | 8.04 |
| PD1<sub>n3</sub> DPA | 0.00 | ± | 0.00 | 0.00 | ± | 0.00 | 0.00 | ± | 0.00 | 0.62 | ± | 0.43 | 0.40 | ± | 0.28 | 0.50 | ± | 0.37 |
| 10S,17S-diHDPA | 19.08 | ± | 5.20 | 20.59 | ± | 4.75 | 21.85 | ± | 6.21 | 16.40 | ± | 4.83 | 17.32 | ± | 5.45 | 22.76 | ± | 7.84* |
| MaR1<sub>n3</sub> DPA | 0.00 | ± | 0.00 | 0.00 | ± | 0.00 | 0.00 | ± | 0.00 | 0.00 | ± | 0.00 | 0.00 | ± | 0.00 | 0.00 | ± | 0.00 |
| 7S,14S-diHDPA | 53.09 | ± | 10.49 | 50.59 | ± | 8.68 | 59.71 | ± | 15.52 | 41.39 | ± | 12.98 | 41.42 | ± | 14.33 | 53.73 | ± | 20.22 |
| **EPA bioactive metabolome** |  |  |  |  |  |  |  |  |  |  |  |  |  |  |  |  |  |  |
| RvE1 | 0.00 | ± | 0.00 | 0.00 | ± | 0.00 | 1397.39 | ± | 315.54 | 0.00 | ± | 0.00 | 0.00 | ± | 0.00 | 1306.85 | ± | 325.88 |
| RvE2 | 33.49 | ± | 1.82 | 32.39 | ± | 1.80 | 31.46 | ± | 2.60 | 37.50 | ± | 1.88 | 32.74 | ± | 3.15* | 37.98 | ± | 2.17 |
| RvE3 | 3.12 | ± | 0.67 | 3.46 | ± | 0.41 | 3.13 | ± | 0.67 | 2.58 | ± | 0.55 | 2.59 | ± | 0.97 | 3.29 | ± | 1.10 |
| **AA bioactive metabolome** |  |  |  |  |  |  |  |  |  |  |  |  |  |  |  |  |  |  |
| LXA<sub>4</sub> | 2.02 | ± | 0.16 | 2.39 | ± | 0.30 | 2.15 | ± | 0.22 | 1.68 | ± | 0.26 | 1.75 | ± | 0.20 | 1.42 | ± | 0.19 |
| LXB<sub>4</sub> | 3.84 | ± | 1.85 | 396.95 | ± | 90.14* | 3.43 | ± | 1.88 | 0.00 | ± | 0.00 | 330.76 | ± | 102.83 | 0.00 | ± | 0.00 |
| 5,15-diHETE | 2332.88 | ± | 550.67 | 2496.44 | ± | 629.07 | 2509.74 | ± | 681.76 | 1860.42 | ± | 589.74 | 1967.32 | ± | 667.77 | 2461.83 | ± | 908.88 |
| 15-epi-LXA<sub>4</sub> | 56.73 | ± | 7.92 | 50.34 | ± | 6.70* | 49.89 | ± | 9.07* | 54.85 | ± | 9.78 | 51.37 | ± | 10.54 | 46.96 | ± | 7.07* |
| 15-epi-LXB<sub>4</sub> | 12.52 | ± | 2.10 | 13.37 | ± | 1.08 | 12.82 | ± | 1.80 | 11.42 | ± | 1.74 | 10.90 | ± | 1.87 | 13.05 | ± | 1.68* |
| 13,14-dihydro-15-oxo-LXA<sub>4</sub> | 28.45 | ± | 4.08 | 23.87 | ± | 4.04* | 22.27 | ± | 3.65* | 29.75 | ± | 3.14 | 25.87 | ± | 3.29 | 25.64 | ± | 1.23 |
| 15-oxo-LXA<sub>4</sub> | 22.42 | ± | 15.50 | 18.62 | ± | 12.75 | 22.04 | ± | 15.10 | 25.43 | ± | 17.48 | 22.32 | ± | 15.31 | 18.91 | ± | 13.61 |
| LTB<sub>4</sub> | 0.00 | ± | 0.00 | 0.00 | ± | 0.00 | 0.00 | ± | 0.00 | 0.00 | ± | 0.00 | 0.00 | ± | 0.00 | 0.00 | ± | 0.00 |
| 5,12 diHETE | 333.87 | ± | 61.15 | 324.79 | ± | 57.51 | 332.24 | ± | 58.63 | 261.78 | ± | 76.72 | 260.65 | ± | 77.52 | 277.98 | ± | 90.72 |
| 6-trans-LTB<sub>4</sub> | 133.21 | ± | 12.31 | 124.72 | ± | 14.20 | 126.85 | ± | 12.30 | 134.89 | ± | 29.45 | 131.64 | ± | 28.06 | 139.94 | ± | 32.82 |
| 12-epi-6-trans-LTB<sub>4</sub> | 164.92 | ± | 28.04 | 146.95 | ± | 25.43 | 149.35 | ± | 23.89 | 152.83 | ± | 42.66 | 150.40 | ± | 40.20 | 164.86 | ± | 49.33 |
| PGD<sub>2</sub> | 359.4<br>2 | ± | 160.2<br>3 | 297.7<br>1 | ± | 83.52 | 327.9<br>5 | ± | 114.5<br>6 | 386.2<br>8 | ± | 91.64 | 348.8<br>6 | ± | 73.43 | 332.9<br>8 | ± | 61.52 |
| PGE<sub>2</sub> | 6475.<br>47 | ± | 4794.<br>74 | 6574.<br>62 | ± | 4996.<br>22 | 6342.<br>04 | ± | 4649.<br>23 | 8854.<br>74 | ± | 3776.<br>51 | 9020.<br>79 | ± | 4111.<br>85 | 9503.<br>78 | ± | 3911.6<br>5* |
| PGF<sub>2α</sub> | 273.1<br>7 | ± | 227.8<br>6 | 227.4<br>5 | ± | 191.6<br>1 | 221.5<br>4 | ± | 179.6<br>5 | 328.7<br>8 | ± | 184.2<br>9 | 277.5<br>8 | ± | 176.0<br>2 | 247.9<br>5 | ± | 168.56<br>* |
| TXB<sub>2</sub> | 10.20 | ± | 0.84 | 9.21 | ± | 0.71* | 9.19 | ± | 0.52 | 14.32 | ± | 3.36 | 13.92 | ± | 1.87 | 11.78 | ± | 1.32 |

Tendon stromal cells (60,000 cells per well) were derived from patients with shoulder tendon tears (Diseased n=5 donors) or healthy volunteer hamstring tendons (Healthy n=5 donors) and incubated in the presence of 10nM LXB<sub>4</sub>, 10nM RvE1 or vehicle only in the presence of 10ngml<sup>-1</sup> IL1β. Cell incubations were terminated using ice-cold methanol containing internal standards and lipid mediators (LM) were identified and quantified using LM-profiling (see methods for details). Results are expressed as pg/incubation. Mean ± SEM of n = 5 per incubation. \* p ≤ 0.05 comparison between respective vehicle incubated tendon stromal cells. The detection limit was ~ 0.1 pg. 0.0, Below limits of detection.

