## Supplementary figures and images for "Proresolving mediators LXB_4_ and RvE1 regulate inflammation in stromal cells from patients with shoulder tendon tears"

### Figure S1

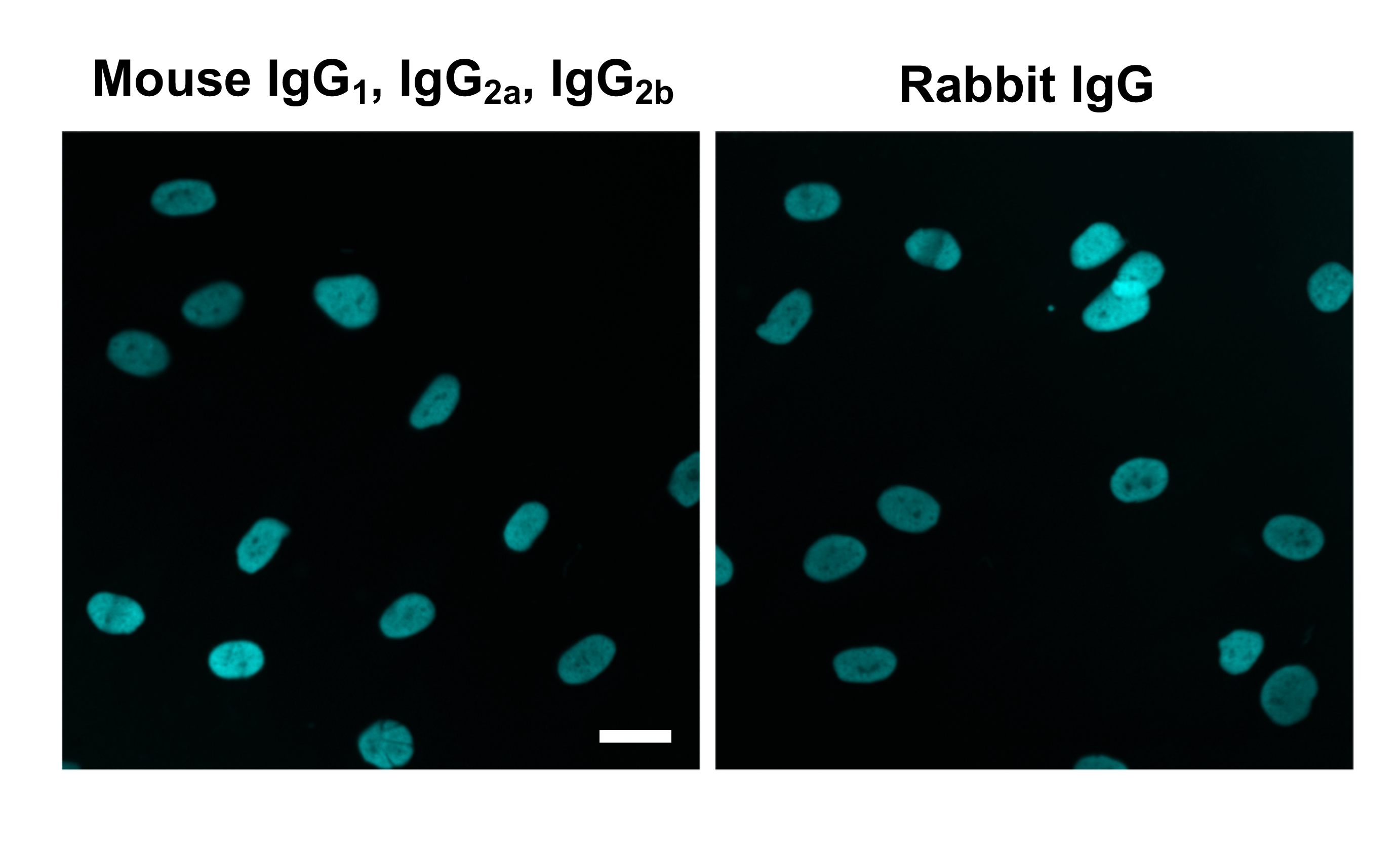
